## Supplementary material for "Bacterial dormancy: a subpopulation of viable but non-culturable cells demonstrates better fitness for revival": S1

**S1, Results: Factors to enable VBNC formation**

**Results:** Initial experiments showed that we could generate a microcosm in which cells entered a VBNC state and where we could resuscitate them to culturable forms depended on several factors including; a) age of strain, b) minimising the damage to cells during preparation, c) conditions that induce the VBNC state and d) conditions provided for resuscitation. ***Age of Strain:*** To establish laboratory microcosms, bacterial stocks were removed from frozen storage and allowed to acclimatise on agar at room temperature for 5 days (fresh cultures) or <14 days (older cultures). Figure 1A shows the response of *V. parahaemolyticus* RIMD2210633 cells subjected to nutritional stress (stored in modified PBS) and cold storage (incubated at 6-9 ^°^C). When fresh cultures (~5 days old) were used to set up the microcosms, plate counts declined to undetectable levels within 30-35 days (Figure 1A). In microcosms set up using older cultures that were stored on agar for more than 2 weeks, plate counts decline faster taking approximately 20 days to reach undetectable levels (Figure 1A). **Microcosm preparation:** During preparation of bacterial cells high-speed centrifugation resulted in the production of a large cell pellet. By allowing the pellet to be incubated in fresh media for 5-10 minutes followed by gentle resuspension with a sterile loop caused less damage to the cells (which we observed by propidium iodide (PI) staining) and increased the likeliness of the cells entering a VBNC state. **Conditions that induce the VBNC state**: We observed that when microcosms were stored at 4 °C, cell-counts declined within a few days to undetectable levels and using PI staining cells were mainly dead or damaged. This was also confirmed by an inability to resuscitate cells from the microcosm. Consequently, the ideal temperature for maintaining the microcosms was established to be between 6 – 9 °C. **Conditions provided for resuscitation:** Within microcosms, the number of culturable cells decreased over time while the number of VBNC cells increased. This suggests that not all the VBNC cells are of the same age and entering the VBNC is a stochastic process. Thus the resuscitation window is defined as the period of time in which VBNC cells can be revived in response to a stimulus under study and may vary for each microcosm. To maintain reproducibility in resuscitation we analysed the VBNC cells in the microcosm when there were no culturable cells growing on routine agar and used PBS as a medium to revive and determine the number of cells in the VBNC state. Our previous experiments showed that *V. parahaemolyticus* can survive in PBS but not actively divide and grow which make it an ideal medium to assess VBNC populations. Cells from microcosms that were unculturable on routine agar but demonstrated a return to growth phenotype when placed in PBS and subjected to a slow increase in temperature where defined as VBNC cells.
