## Supplementary material for "Bacterial dormancy: a subpopulation of viable but non-culturable cells demonstrates better fitness for revival": S2

**S2: Protein data and the numbers of proteins detected in each group.**

| Sample | Number of proteins detected |
| --- | --- |
| T0 | 1533 |
| P1-T12 | 1477 |
| P2-T12 | 1497 |
| P1-T50 | 1444 |
| P2-T50 | 1449 |
