## Supplementary material for "Bacterial dormancy: a subpopulation of viable but non-culturable cells demonstrates better fitness for revival": S4

**S4. Correlation between the proteomes of the analysed groups.** Determined by regression analysis. Mean of the normalised abundance values were used with each group.

| **Sample comparisons** | **Adjusted R^2^ of linear regression** | **Slope of the linear regression line** |
| --- | --- | --- |
| **T0 vs P2-T12** | 0.601 | 0.718 |
| **T0 vs P1-T12** | 0.494 | 0.596 |
| **T0 vs P2-T50** | 0.351 | 0.397 |
| **T0 vs P1-T50** | 0.258 | 0.327 |
| **P2-T12 vs P1-T12** | 0.796 | 0.817 |
| **P2-T50 vs P1-T50** | 0.799 | 0.859 |
| **P2-T12 vs P2-T50** | 0.524 | 0.524 |
| **P1-T12 vs P1-T50** | 0.555 | 0.565 |
