## Supplementary material for "Bacterial dormancy: a subpopulation of viable but non-culturable cells demonstrates better fitness for revival": S5

### S5: Principal component analysis of global protein expression profiles of *V. parahaemolyticus* obtained from bacteria grown in vitro and in VBNC state. Replicates of the Log Phase cells, T0 and T12 samples are clustered together indicating experimental reproducibility. Replicates of the T50 samples are more distant from each other.

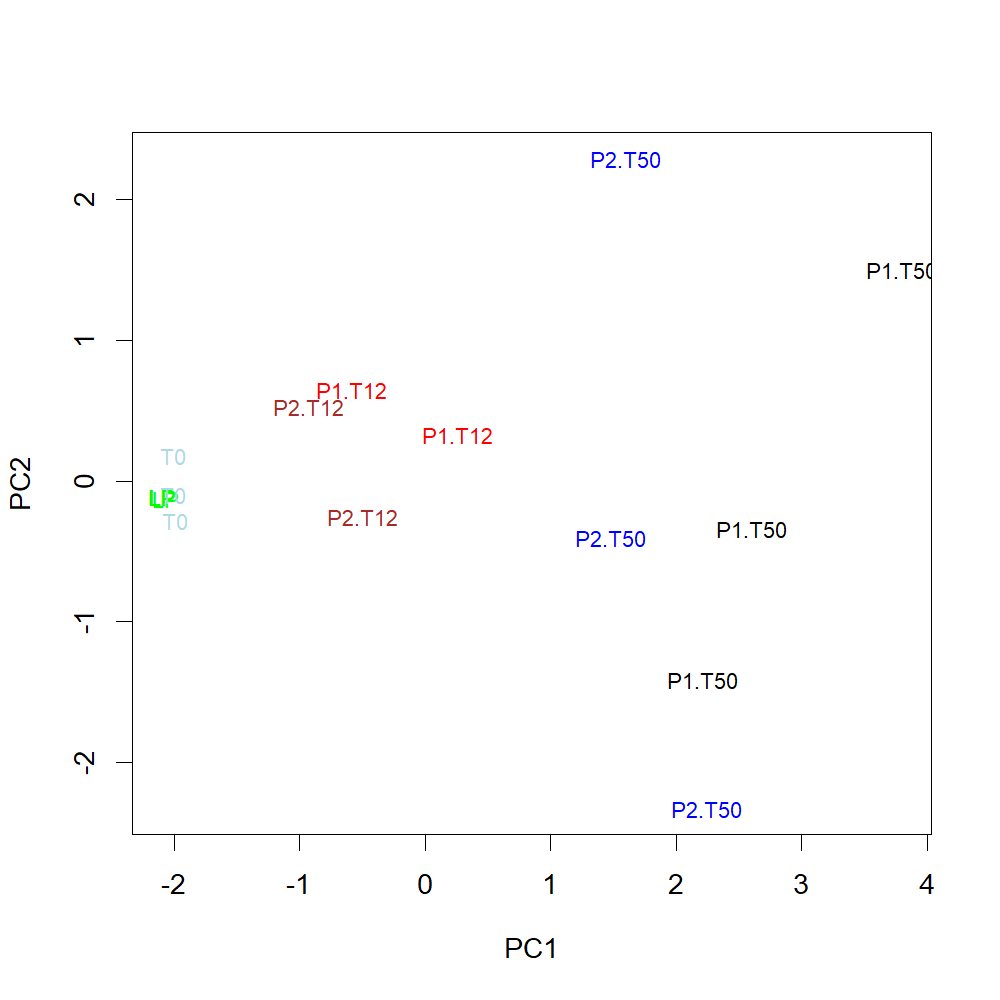
