## Supplementary figures and images for "Bacterial dormancy: a subpopulation of viable but non-culturable cells demonstrates better fitness for revival"

### S7

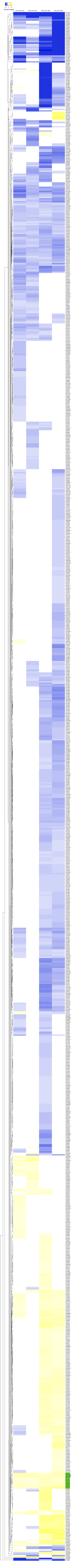
